# Impact of phosphorylation on thermal stability of proteins

**DOI:** 10.1101/2020.01.14.903849

**Authors:** Clément M. Potel, Nils Kurzawa, Isabelle Becher, Athanasios Typas, André Mateus, Mikhail M. Savitski

## Abstract

Reversible protein phosphorylation regulates virtually every cellular process and is arguably the most well-studied post-translational modification. Still, less than 3% of the phosphorylation sites identified in humans have annotated functions. Functionally-relevant phosphorylation sites are known to trigger conformational changes to proteins and/or to regulate their interactions with other proteins, nucleic acids and small molecules - all of which can be reflected in the thermal stability of a protein. Thus, combining thermal proteome profiling (TPP) with phosphoproteomics (phospho-TPP) provides a way to assess the functional relevance of identified phosphorylation sites on a proteome-wide scale by comparing the melting behavior of a protein and its phosphorylated form(s). We performed phospho-TPP experiments in HeLa cells with an optimized protocol, and conclude that phosphorylation does affect protein thermal stability, but to a much lesser extent than previously reported.

## Introduction

Reversible protein phosphorylation regulates virtually every cellular process and is arguably the most well-studied post-translational modification.^1^ Still, less than 3% of the phosphorylation sites identified in humans have annotated functions.^2,3^ Functionally-relevant phosphorylation sites are known to trigger conformational changes to proteins and/or to regulate their interactions with other proteins, nucleic acids and small molecules—all of which can be reflected in the thermal stability of a protein.^4^ Thus, combining thermal proteome profiling (TPP)^5^ with phosphoproteomics^6^ (phospho-TPP) provides a way to assess the functional relevance of identified phosphorylation sites on a proteome-wide scale by comparing the melting behavior of a protein and its phosphorylated form(s).

The first application of phospho-TPP was described by Azimi et al.^7^, though a systematic comparison between protein thermal stabilities and phosphorylated protein forms was not performed. Recently, Huang et al.^8^ investigated the effect of 2,883 phosphorylation sites on protein thermal stability, and reported significant changes in the melting behavior for 719 (25%) of these sites (without multiple testing correction). Moreover, the correlation between the melting point of a protein and its phosphorylated form(s) was surprisingly low (R^2^=0.18; Fig. 1A; Supplementary Fig. 1). This would imply that most phosphorylation events fundamentally reshape the thermal stability of proteins. While this constitutes an exciting hypothesis, we hereby show that this low correlation is likely technically-rather than biologically-driven. Indeed, when assessing the reproducibility of the published protein melting point (T_m_) data,^8^ we found low correlations between replicates, both for the melting points of non-modified (average R^2^ of 0.43 between replicates) (Supplementary Fig. 2) and of phosphorylated proteins (average R^2^ of 0.22) (Supplementary Fig. 3)—despite the claims of high reproducibility of T_m_ by the authors (see Supplementary discussion on ‘T_m_ estimate reproducibility’). After carefully re-evaluating their work, we identified multiple key points that contributed to this poor reproducibility and the unexpected high impact of phosphorylation on protein thermal stability (see Supplementary discussion). Therefore, we performed phospho-TPP experiments in HeLa cells with an optimized protocol (Supplementary Fig. 4), and conclude that phosphorylation does affect protein thermal stability, but to a much lesser extent than previously reported (Fig. 1A-B).^8^ This is reinforced by an accompanying analysis by an independent research group^9^.

**Fig. 1.**
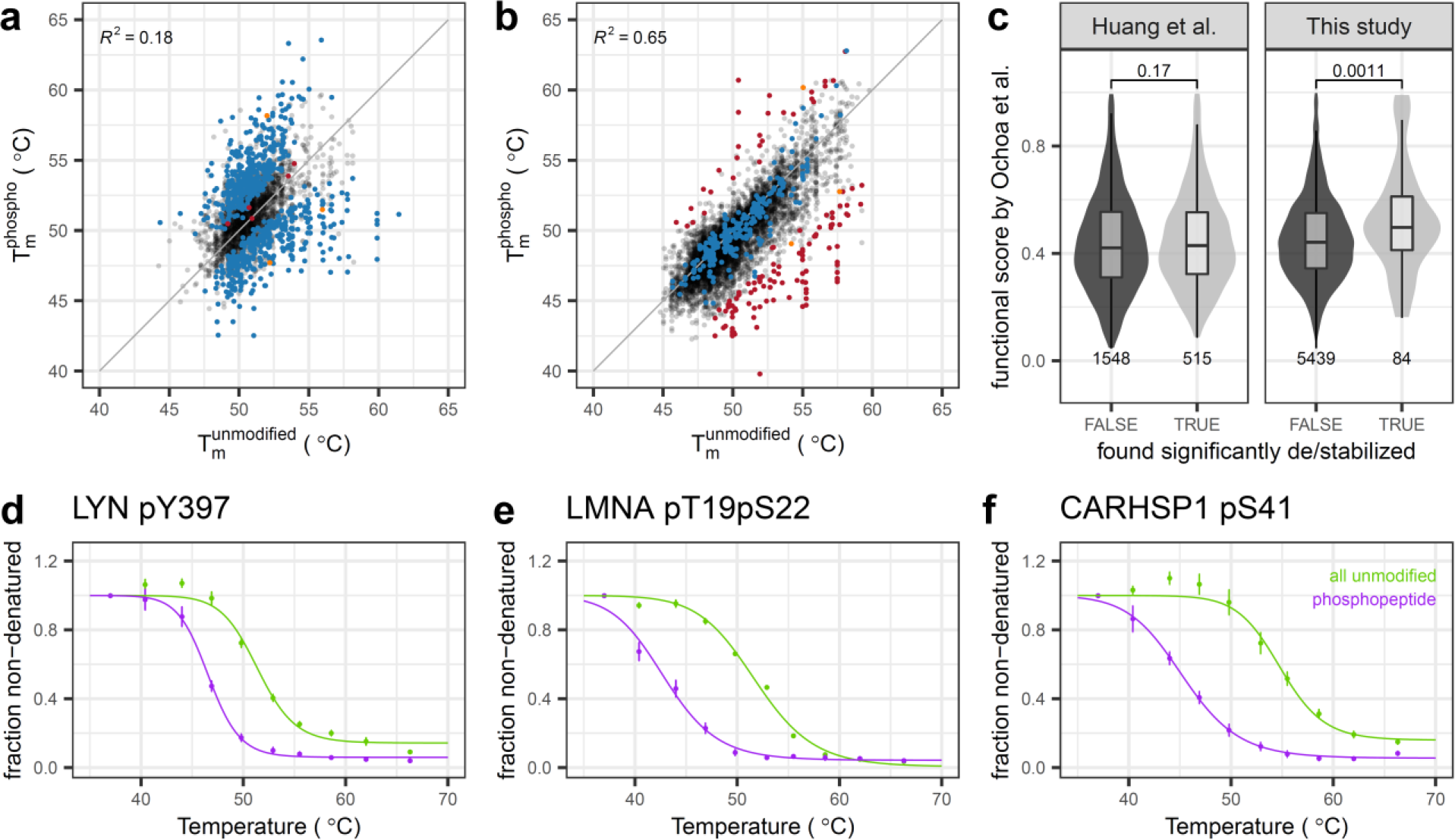
Global impact of phosphorylation on thermal stability of proteins is less extensive than previously reported. (a-b) Comparison of non-modified and phosphorylated protein T_m_ in (a) Huang et al.^8^ and this study (b). The grey line represents the identity line, points are colored according to: hits called significant by the original work (blue), hits found by this study (red) and overlapping hits between the two studies (orange). (c) Comparison of predicted functional importance of significant and non-significant phosphosites between the two studies (Wilcoxon signed-rank test)—the functional score aggregates multiple parameters for each phosphosite using machine learning and ranges from 0 to 1, with a higher value representing a higher probability that the phosphosite has functional relevance^12^. Center line in box plots represents the median, bounds of the boxes are the 25^th^ and 75^th^ percentiles, and the whiskers correspond to the highest or lowest value, or if the lowest or highest value is an outlier (i.e., greater than 1.5-fold of the interquartile range), they correspond to 1.5-fold of the interquartile range. (d-f) Examples of melting curves of significant hits found in this study. Error bars represent standard error of the mean.

## Results and Discussion

Our phospho-TPP analytical strategy resembled the approach by Huang et al.^8^ (Supplementary Fig. 4), but with critical differences in that: i) we labeled peptides for quantification with tandem mass tags (TMT) *prior* to phosphopeptides enrichment to eliminate potential biases introduced by the individual phosphopeptide enrichment of the different temperatures and the separate labeling of non-modified and phosphopeptides; ii) we used two orthogonal chromatographic separations prior to mass spectrometry analysis to gain experimental depth and minimize TMT ratio compression arising from peptides co-isolation;^9^ and iii) we implemented a robust data analysis approach including two data normalization steps and multiple testing correction (Supplementary table). In total, we identified 42,051 unique phosphopeptides across five biological replicates (Supplementary data 1-2). Notably, phosphopeptides constituted 98% of reliably identified peptides in the enriched samples—contrary to Huang et al.^8^, in which a reanalysis of their data yielded 61% of phosphopeptides (noted to be variable between replicates in an accompanying Matters Arising^9^). Our approach yielded high T_m_ reproducibility between biological replicates of both non-modified (average R^2^ of 0.86; Supplementary Fig. 5) and phosphorylated proteins (average R^2^ of 0.78; Supplementary Fig. 6). Further, we compared the T_m_ of non-modified proteins to previously published datasets^10,11^ (including from other cell lines and acquired in different laboratories; Supplementary Fig. 7), and observed substantially higher correlation with our dataset (average R^2^ of 0.75) than with Huang et al.^8^ (average R^2^ of 0.43).

With our approach, we could confidently compare the melting behavior of 7,864 unique phosphopeptides with their corresponding proteins (these represented cases for which we had data for the phophopeptide and the corresponding non-modified protein in at least three replicates, and a high quality melting curve; Supplementary data 3). In contrast to Huang et al.^8^ (Fig. 1A), we observed a good agreement between the T_m_ of proteins and phosphorylated form(s) (R^2^=0.65; Fig. 1B), with 129 phosphopeptides showing a statistically significant difference in their melting behavior compared to their respective proteins. These phosphorylation sites showed significantly higher predicted functional relevance^12^ (p = 0.0011, Wilcoxon signed-rank test; Fig. 1C), suggesting that sites with altered thermal stability are likely to be biologically relevant. This was not the case for the significant sites annotated by Huang et al.^8^ (p = 0.17; Fig. 1C). Even though we identified significant differences in melting behavior of phosphorylation sites for some of the examples highlighted in Huang et al.^8^ (e.g., pSer58 of TPI1, which is thermally destabilized), these cases constitute the exception rather than the rule. This was further emphasized by the fact that from the 237 significant phosphosites identified in Huang et al.^8^ that were among our 7,864 high quality cases, the majority of them (98.7%; 234 sites) showed no significant change in melting behavior (Fig. 1B). Taken together, our study shows that the structural remodeling due to phosphorylation is not as widespread as previously reported, and that overall there is a good agreement between the melting behavior of proteins and phosphorylated proteoforms.

Despite the general concordance of melting behaviors of phosphorylated and non-phosphorylated forms of proteins, we observed a number of cases where phosphorylation significantly impacted the thermal stability of proteins (Fig. 1D-F). For example, phosphorylation on Tyr397 of the non-receptor tyrosine-protein kinase Lyn, which is involved in the regulation of immune response, hematopoiesis and response to DNA damage,^13^ led to a significant decrease in thermal stability of this protein (ΔT_m_ = −5.2 °C; Fig. 1D). Phosphorylation on this site within the activation loop of Lyn influences its activity by inducing a known conformational change that increases accessibility to the ATP binding pocket.^13^ Another interesting example are the phosphorylations on Thr19 and Ser22 of Lamin A, which led to a significant decrease of thermal stability (ΔT_m_ = −8.6 °C; Fig. 1E). Lamin A is a nuclear intermediate filament protein that regulates chromatin organization, replication and transcription through DNA and transcription factor binding,^14^ with phosphorylation of Thr19 and Ser22 being essential for its disassembly during mitosis.^15^ Finally, the calcium-regulated heat-stable protein CARHSP1 contains a cold-shock domain that binds RNA and single-stranded DNA, which stabilizes the tumor necrosis factor mRNA in processing bodies and exosomes.^16^ In accordance with the fact that a Ser41 phosphomimetic mutant decreases the affinity towards nucleic acids and abolishes protein localization in stress granules,^17^ we detected a significant decrease in protein thermal stability for the phosphorylated Ser41 proteoform (ΔT_m_ = −9.3 °C; Fig. 1F). In this case, phospho-TPP was able to resolve the thermal stability effect of Ser41 phosphorylation from that of adjacent phosphosites in the same phosphopeptide, for which melting behavior resembled that of the non-modified protein (Supplementary Fig. 8).

In conclusion, we present a strategy for performing and analyzing phospho-TPP experiments and provide a high quality dataset, which will serve as a resource for understanding the effects of phosphorylation on protein thermal stability. Phosphorylation sites which have a significant effect on protein thermal stability are more likely to be functionally relevant, by modulating protein conformation, localization, and protein interactions.

## Acknowledgements

This work was supported by the European Molecular Biology Laboratory. C.P. and A.M. were supported by a fellowship from the EMBL Interdisciplinary Postdoc (EI3POD) programme under Marie Skłodowska-Curie Actions COFUND (grant number 664726). N.K. was supported by a fellowship of the EMBL International PhD programme (EIPP).

## Author contributions

C.P., I.B., A.M. performed the experimental work. N.K. performed the data analysis. All authors contributed to the interpretation of the data and writing of the manuscript.

## Supplementary information

### Methods

#### Cell culture

HeLa Kyoto cells were cultured at 37 °C, 5% CO_2_, in DMEM including 1 mg/ml glucose, supplemented with 10% fetal bovine serum and 1 mM glutamine.

#### Thermal proteome profiling

Thermal proteome profiling was performed as previously described.^1–3^ Briefly, for each replicate ten 2×10^7^ HeLaK aliquots were heated for 3 min to a range of temperatures (37.0-40.4-44.0-46.9-49.8-52.9-55.5-58.6-62.0-66.3 °C). After another 3 min incubation at room temperature, cells were lysed for 1 h at 4 °C (lysis buffer: 0.8 % NP-40, 1.5 mM MgCl_2_, cOmplete protease inhibitors, PhosStop, benzonase, 2 mM NaF, 2 mM Na_3_VO_4_, 2 mM Na_4_O_2_P_7_ in PBS). Heat-induced protein aggregates were removed by filter plates (Millipore, 0.45µm) and the soluble fraction was used for protein concentration determination.

#### Protein digestion and peptide labelling

Approximately 300 µg protein (determined for the lowest temperature) were diluted with sonication buffer (1% sodium deoxycholate, 5 mM tris(2-carboxyethyl)phosphine, 30 mM chloroacetamide, 1 mM MgCl_2_, 1% benzonase) and sonicated in bioruptor for 15 cycles (30 s on/30 s off). Another incubation with 1% benzonase was performed for at least 30 min at room temperature, to remove nucleic acids, which are common contaminants of phosphopeptide enrichment.^4^

Proteins were digested according to a modified SP3 protocol.^5,6^ Briefly, protein samples were added to the bead suspension (Thermo Fisher Scientific—Sera‐Mag Speed Beads, CAT# 4515‐2105‐050250, 6515‐ 2105‐050250) in ethanol. After a 15 min incubation at room temperature with shaking, beads were washed four times with 70% ethanol. Next, proteins were digested overnight by adding 100 μl of digest solution (30 mM chloroacetamide, 5 mM TCEP, 1 µg/µl trypsin, in 100 mM HEPES pH 8). Peptides then were eluted from the beads, dried under vacuum, reconstituted in 20 μl of water, and labeled for 60 min at room temperature with 800 µg of TMT10plex (Thermo Fisher Scientific) dissolved in 8 μl of acetonitrile. The reaction was quenched with 8 μl of 5% hydroxylamine, and experiments belonging to one TPP-TR replicate were combined. Samples were acidified with trifluoroacetic acid (TFA, final concentration of 1%) and desalted with solid‐phase extraction by loading the samples onto a Waters t-C18 SepPak 50 mg column, washing them twice with 1 ml of 0.1% TFA, eluting them with 400 μl of 50% acetonitrile acidified with 0.1% TFA, before lyophilisation.

#### Phosphopeptide enrichment

The phosphopeptide enrichment was performed generally as previously described^4^. Briefly, lyophilized peptides were resuspended in buffer A (70% ACN, 0.07% TFA) and injected on a ProPac IMAC-10 column (Thermo Fisher Scientific, 4×50 mm) loaded with Fe^3+^ cations. Peptide loading was performed at a flow rate of 400 µL/min for 6 minutes using an Ultimate 3000 UHPLC liquid chromatography system (Thermo Fisher Scientific). After washing with 100% buffer A for 6 minutes at 1 ml/min, phosphopeptides were eluted by switching to 50% buffer B (0.3% ammonia) for 2 minutes, at a flow rate of 500 µl/min. Both the non-bound and phosphopeptides fractions were collected before lyophilisation.

#### High pH fractionation

An aliquot (5%) of the non-bound fraction from the phosphopeptide enrichment step was fractionated onto 29 fractions on a Phenomenex Gemini 3 µm C18 110 Å 100 mm x 1 mm column under high pH conditions.^5^ This consisted of an 85 min gradient (mobile phase A: 20 mM ammonium formate (pH 10) and mobile phase B: acetonitrile) at a 0.1 ml/min starting at 0% B, followed by a linear increase to 35% B from 2 min to 60 min, with a subsequent increase to 85% B from up to 62 min and holding this up to 68 min, which was followed by a linear decrease to 0% B up to 70 min, finishing with a hold at this level until the end of the run. Fractions were collected every two minutes from 12 min to 70 min and every 12^th^ fraction was pooled together.

Phosphopeptides were fractionated using in-house packed C18 microcolumns. To do so, gel loader tips were plugged with C18 resin (Affinisep AttractSPE C18 Disks) before packing with 1 mg of C18 material (Dr Maisch, 5 µm, 120 Å). Dried phosphopeptides were resuspended in 40 µL buffer A (20 mM ammonium formate at pH 10) before loading onto the microcolumn by centrifugation (loading speed ≈ 10 µl/min) and wash with 10 µL buffer A. Phosphopeptides were fractionated using a stepped gradient of acetonitrile consisting of sequential addition/elutions with 10 µl of the following solutions: 1%, 3%, 5%, 7%, 9%, 11%, 13%, 15%, 17%, 19%, 21%, 23%, 24%, 26%, 28%, 30%, 35%, 40% acetonitrile/buffer A (elution speed ≈ 10 µl/min). The flow-through and wash were collected and correspond to the fraction FT, while elutions with 1% and 3% acetonitrile were pooled and correspond to fraction F0. The other elutions were pooled as follow: F1 = 5%, 17%, 28%; F2 = 7%, 19%, 30%; F3 = 9%, 21%, 35%; F4 = 11%, 23%, 40%; F5 = 13%, 24%; F6 = 15%, 26%.

#### LC-MS/MS measurements

Peptides were separated using an UltiMate 3000 RSLCnano system (Thermo Fisher Scientific) equipped with a trapping cartridge (Precolumn; C18 PepMap 100, 5 μm, 300 μm i.d. × 5 mm, 100 Å) and an analytical column (Waters nanoEase HSS C18 T3, 75 μm × 25 cm, 1.8 μm, 100 Å). Solvent A was 0.1% formic acid in LC‐MS grade water and solvent B was 0.1% formic acid in LC‐MS grade acetonitrile. Peptides were loaded onto the trapping cartridge (30 μl/min of solvent A for 3 min) and eluted with a constant flow of 300 nl/min using a 120 min analysis time (with a 10–28% B linear gradient, followed by an increase to 40% B, and re‐equilibration to initial conditions). The LC system was coupled to a Fusion Lumos Tribrid mass spectrometer (Thermo Fisher Scientific) operated in positive ion mode with a spray voltage of 2.2 kV and capillary temperature of 275°C. Full‐scan MS spectra with a mass range of 375–1,500 m/z were acquired in profile mode in the Orbitrap using a resolution of 120,000 (maximum injection time of 50 ms and automatic gain control (AGC) set to 4e5 charges). The mass spectrometer was operated in data dependent acquisition mode with a maximum duty cycle time of 3 s. The most intense precursors with charge states 2–7 and a minimum intensity of 2e5 were selected for subsequent HCD fragmentation (isolation window of 0.7 m/z and normalized collision energy of 36%) with a 60 seconds dynamic exclusion window, and MS/MS spectra were acquired in profile mode with a resolution of 30,000 in the Orbitrap (maximum injection time of 94 ms and AGC target of 1e5 charges).

Phosphopeptide fractions were analyzed on the same LC-MS/MS system with a few differences listed below. Phosphopeptides were resuspended in a mixture of 50 mM citric acid and 1% formic acid before loading, trapping and separation using a linear gradient from 8% to 25% buffer B, followed by an increase to 40% buffer B (total analysis time of 120 min). Full scans were acquired in the Orbitrap with a scan range of 375–1,400 m/z, and precursors were sequentially isolated and fragmented with a 30 seconds dynamic exclusion window. MS/MS spectra were acquired in the Orbitrap at a resolution of 30,000 with an AGC target of 1e5 charges and a maximum injection time of 110 ms.

#### Peptide and protein identification

Mass spectrometry data were processed with isobarQuant,^2,7^ and the identification of peptide and protein was performed with Mascot 2.4 (Matrix Science) against the human UniProt FASTA (Proteome ID: UP000005640), modified to include known contaminants and the reversed protein sequences (search parameters: trypsin; missed cleavages 3; peptide tolerance 10 ppm; MS/MS tolerance 0.02 Da; fixed modifications were carbamidomethyl on cysteines and TMT10plex on lysine; variable modifications included acetylation on protein N‐terminus, oxidation of methionine, and TMT10plex on peptide N‐termini).

In parallel, the phosphopeptides-enriched raw data files were processed by the MaxQuant software^8^ (version 1.6.2.3) to assess phosphorylation sites localization probabilities. Files were searched against a reviewed *Homo sapiens* database (UniProt, September 2018), with the following parameters: trypsin digestion (cleavage at the C-term of lysine and arginine, even when followed by a proline residue) with a maximum of 3 missed cleavages, TMT10plex labeling, fixed carbamidomethylation of cysteines, variable oxidation of methionines, as well as, variable phosphorylation of serine, threonine and tyrosine residues. Mass tolerance was set to 4.5 ppm at the MS1 level and 20 ppm at the MS2 level. A score cut-off of 40 was used for modified peptides, the false discovery rate was set to 0.01 and the minimum peptide length to 7 residues.

#### Data preprocessing

Search results for phosphopeptide-enriched samples from isobarQuant^2,7^ and MaxQuant^8^ were merged based on peptide MS/MS scan ID and kept only for further analysis when they had at least one phosphosite which could be localized with a probability higher than 0.75 (extracted from MaxQuant output)—to only include phosphosites with high localization confidence, generally termed class I phosphosites^9^—, signal to interference ratio (S2I) equal or higher than 0.5 and precursor to threshold ratio (P2T) equal or higher than 4 (both extracted from isobarQuant output)—to minimize ratio compression originating from co-isolated peptides^1^. Peptides with identical sequence and phosphorylation localization were merged by summing their signal intensities. Search results for the non-modified fraction were used only based on isobarQuant output parameters requesting that peptides were unique for the protein, a Mascot score of higher than 20 and an FDR lower than 0.01. Measured signals for each unique peptide in each replicate and condition were transformed into fold-changes by dividing signal intensities by each corresponding measurement at the lowest temperature (37 °C).

#### Data normalization

To normalize the fold-changes of each phosphopeptide enriched sample to each respective non-modified sample, we retrieved normalization-factors to align median fold changes at each temperature of jointly identified non-modified peptides. This was implemented by finding overlapping non-modified peptides between matching replicates of phosphopeptide enriched and non-modified samples fulfilling the following filter criteria: 7^th^ temperature should be between 0.3 and 0.7, fold change at 9^th^ temperature should be between 0 and 0.4, and fold change at 10^th^ temperature should be between 0 and 0.2. Then, normalization-factors were determined for fold changes at each temperature by computing the ratio between the median fold change of these peptides in the non-modified and phosphopeptide enriched samples. Finally, retrieved normalization-factors were applied to normalize the fold changes at each measured temperature of all peptides within the phosphopeptide enriched sample.

Next, we applied a previously described curve-based normalization procedure between replicates of the non-bound fraction samples.^10^ Briefly, this involved: i) building the set of overlapping peptides between all datasets; ii) filtering these peptides to fulfill the following criteria: fold change measured at 3^rd^ temperature should be higher than 1, fold change at 7^th^ temperature should be between 0.3 and 0.7, fold change at 9^th^ temperature should be between 0 and 0.4, and fold change at 10^th^ temperature should be between 0 and 0.2; iii) finding the largest subset of peptides fulfilling these criteria in either of the replicates; iv) fitting a melting curve to the median fold change of the set of selected peptides for each replicate; v) the replicate yielding the highest R^2^ value was selected and normalization factors for each of the replicates were obtained by computing the fractions of fold changes predicted by that melting curve at each given temperature and the median fold changes measured for each respective replicate; vi) retrieved normalization factors were then applied to both the non-modified fraction and the respective phosphopeptide-enriched replicates.

#### Melting curve fitting

Melting curves were fitted to each replicate of each phosphopeptide separately using the *drc* R package^11^ with the sigmoidal *LL.4* model:

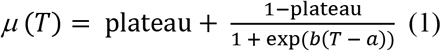

Where *T* is the temperature, plateau is the lower bound, *b* is the slope and *a* the inflection point. The melting point (T_m_) was estimated from the temperature at which *µ(T)* = 0.5, from curves for which R^2^ >0.8 and plateau <0.2.

For the non-modified fraction all peptides mapping to the same protein were jointly fitted per replicate using (1).

#### Detecting differentially melting phosphopeptides

Assessment of differential melting of phosphopeptides compared to all peptides of a given protein in the non-bound fraction was based on inferred melting point (T_m_) comparisons. As previously described,^2^ ΔT_m_ were compared using z-tests and retrieved p-values were adjusted for multiple testing using the method of Benjamini and Hochberg.^12^ Phosphopeptides were accepted as significant hits, when at least two replicates were found with an adjusted p-value < 0.1 and all ΔT_m_s (including non-significant) had consistent signs.

#### Data and code availability

The mass spectrometry proteomics data have been deposited to the ProteomeXchange Consortium via the PRIDE partner repository with the dataset identifier PXD015993. All code to reproduce the analysis is available at https://github.com/nkurzaw/phosphoTPP.

## Supplementary discussion

### T_m_ estimate reproducibility

Huang et al.^13^ assessed data reproducibility by comparing the average T_m_ from biological replicates 1-3 to the average T_m_ from biological replicates 4-6 of the non-modified protein samples (original Supplementary figure 2 in Huang et al.^13^), which yielded a coefficient of determination (R^2^) of 0.7. However, when determining the reproducibility of individual replicates, using the original data from Supplementary Table 1 and 2 in Huang et al.^13^, we found poor correlation between all replicates—both biological (average R^2^ for phosphorylated peptides = 0.21 and average R^2^ for non-modified proteins = 0.42) and technical (average R^2^ for phosphorylated peptides = 0.36 and average R^2^ for non-modified proteins = 0.62).

We have previously shown,10 and reconfirmed here, that T_m_ estimates for non-modified proteins are reproducible (average R^2^ of 0.86; Supplementary Fig. 5), even when considering different studies and different laboratories (R^2^ = 0.82 (same cell line in our lab^14^); R^2^ = 0.69 (different cell line in different lab^15^); Supplementary figure 7). Here, we also observed that the same is true for phosphorylated peptides (Supplementary Figure 6).

Therefore, we showed that protein and peptide T_m_ estimation is reproducible, and that the lack of reproducibility in Huang et al.^13^ might influence their results. The reasons for this poor reproducibility are explored in this work, and we provide a complete protocol and data analysis pipeline to achieve high quality data.

### Peptide labeling and phosphopeptide enrichment

In the work of Huang et al.^4^, the authors, after digesting the soluble protein fraction, saved an aliquot of peptides from each temperature (5% of sample) to estimate the T_m_ of the non-modified proteins. With the remaining 95% of peptide samples, the authors performed separate phosphopeptide enrichments using TiO_2_ tips for each temperature. Two main problems might arise from this approach: i) the separate isobaric labeling of the non-modified and corresponding phosphorylated peptides from the same replicate might introduce unnecessary variability—however, tandem mass tags (TMT) labeling, when the TMT reagent is in stoichiometric excess, generally yields reproducible results,^16^ and therefore this might not be the major contributor to the poor reproducibility of the Huang et al. dataset; ii) the separate phosphopeptide enrichments using TiO_2_ tips for each temperature could introduce larger errors. In quantitative phosphoproteomics, enrichment protocols generally suggest that equal amounts of peptides are used for each conditions. In thermal proteome profiling (TPP) experiments, there are large differences in peptide amounts, in general in the order of 10-fold between the lowest temperature (high peptide amount) and the highest temperature (low peptide amount), which will translate in large differences in peptide input/TiO_2_ ratio between the different temperatures—since the amount of TiO_2_ packed in the tips is not adjusted for each sample. It has been shown that the ratio between peptide and stationary phase is critical to achieve good enrichment selectivity,^17^ and the poor enrichment selectivity observed in the Huang et al. work (phosphopeptides representing roughly 60% of the total peptides identified, while nowadays more than 90% specificity is routinely achieved—98% in our study) is thus an indication that the way the authors performed phosphopeptide enrichment is suboptimal. Further, the enrichment selectivity of Huang et al. was variable from sample to sample, as analyzed in an accompanying work by Smith et al. ^18^. Importantly, the authors did not validate the linearity of enrichment capacity over a broad range of peptide amount input, and therefore this step might negatively influence their results by introducing quantification biases which will impact the estimation of phosphopeptide melting points—particularly, for an approach with low amount of TiO_2_ stationary phase (TiO_2_ spin tips used by the authors, catalog no. 88303, Thermo Fisher Scientific). This lack of linearity of enrichment capacity over a broad range of peptide input amount was previously observed in the case of enrichment performed by Fe^3+^-IMAC in tip format^19,20^, but the same remains to be checked for the TiO_2_ tips used in the Huang et al. study.

All in all, we suggest that a single round of TMT labeling should be performed, and that phosphorylated peptides should be enriched once pooled together. The major disadvantage of this approach is the large amount of TMT reagents needed, since phosphorylated protein forms are generally present at low amounts. However, if a linear enrichment procedure is used, it might be possible to perform TMT labeling post-enrichment.

### Mass spectrometry analysis

Huang et al.^13^ quantified their peptides by analyzing them directly after labeling on an Orbitrap Q-Exactive HF mass spectrometer coupled to liquid chromatography (LC). By exploring the raw data deposited in the MassIVE repository (identifier: MSV000083786), we observed that their effective gradient time is generally lower than 2 h (total analysis time of 2.5 h to 4.3h depending on the replicates) and that an isolation window of 1.2 m/z was used. The high complexity of their samples suggests that the gradients are too short to properly separate peptides and avoid co-isolation during MS2 scans—which is critical for TMT based quantification as peptide co-fragmentation leads to TMT ratio compression.^1^ Indeed, we reprocessed the original data using the MaxQuant software (as described in the methods section), and observed low Parent Ion Fraction (PIF, which represents the ratio of the intensity of selected precursor ions over the summed intensities of all ions present in the isolation window) for the majority of both phosphorylated (median [interquartile range]: 0.78 [0.63-0.90]) and non-modified peptides (0.77 [0.62-0.90]). Our samples, which were pre-fractionated using high pH LC prior to the LC gradient in front of the mass spectrometer, yielding an effective gradient time higher than 20 h (total gradient time of 24 h), resulted in a high PIF (phosphorylated 0.90 [0.78-0.96] and non-modified 0.90 [0.78-0.97] peptides) (Supplementary figure 9). It should be noted that we also used a narrower isolation window of 0.7 m/z to further reduce co-isolation. The high co-isolation observed by Huang et al.13 led to a poor reproducibility of T_m_ estimates (Supplementary Figs. 2–3), a compressed distribution of T_m_ for non-modified proteins (Supplementary Fig. 7) and a poor correlation between T_m_ of non-modified and phosphorylated proteoforms, which biased all the downstream analyses by the authors—since this caused T_m_ comparisons between phosphorylated and non-phosphorylated proteoforms to deviate from the unity line and appear as significant hits (Fig. 1A). This is further highlighted by the narrower distribution of differences in T_m_ between proteins and corresponding phosphorylated proteoforms in our experiments (standard deviation of 1.53 °C) compared to Huang et al.^13^ (standard deviation 2.37 °C; Supplementary Fig. 9).

Overall, we recommend that complex TMT labelled samples be pre-fractionated to reduce sample complexity and avoid ratio compression.^1^

## Supplementary data description

**Supplementary data 1. Phosphopeptide fold-changes at each temperature in each replicate.** (gene_name: Gene name; protein_id: Uniprot ID; sequence: peptide sequence; mod_sequence: modified peptide sequence; phospho_site_STY: number of modified aminoacids in the peptide sequence; rel_fc_TMT_REP_TEMP: relative fold-changes to 37°C (TMT: TMT label; REP: replicate; TEMP: temperature).

**Supplementary data 2. Non-modified peptide fold-changes at each temperature in each replicate.** (gene_name: Gene name; protein_id: Uniprot ID; sequence: peptide sequence; modifications: peptide modifications; rel_fc_TMT_REP_TEMP: relative fold-changes to 37°C (TMT: TMT label; REP: replicate; TEMP: temperature).

**Supplementary data 3. Comparison of melting behavior of phosphopeptide and non-modified peptides.** (gene_name: Gene name; Gene_pSite: Phosphosite position within protein; mod_sequence: modified peptide sequence; mean_phospho: average melting temperature of phosphopeptide; mean_nbf: average melting temperature of non-modified protein; mean_delta_meltPoint: average difference of melting temperature of phosphopeptide and non-modified protein; significant: significantly different melting temperature of phosphopeptide compared to non-modified protein; functional_score: predicted functional importance of phosphosite position, by aggregating multiple parameters for each phosphosite using machine learning and ranges from 0 to 1, with a higher value representing a higher probability that the phosphosite has functional relevance; Tm_phospho_REP: melting temperature of phosphopeptide (REP: replicate); Tm_nbf_REP: melting temperature of non-modified protein (REP: replicate); delta_meltPoint_REP: difference of melting temperature of phosphopeptide and non-modified protein (REP: replicate); p_adj_REP: p-value adjusted for multiple testing correction (REP: replicate); phospho_median_fc_TEMP: median relative fold-changes to 37°C of phosphopeptide (TEMP: temperature); nbf_median_fc_TEMP median relative fold-changes to 37°C of non-modified protein (TEMP: temperature).

## Supplementary figures

**Supplementary figure 1.**
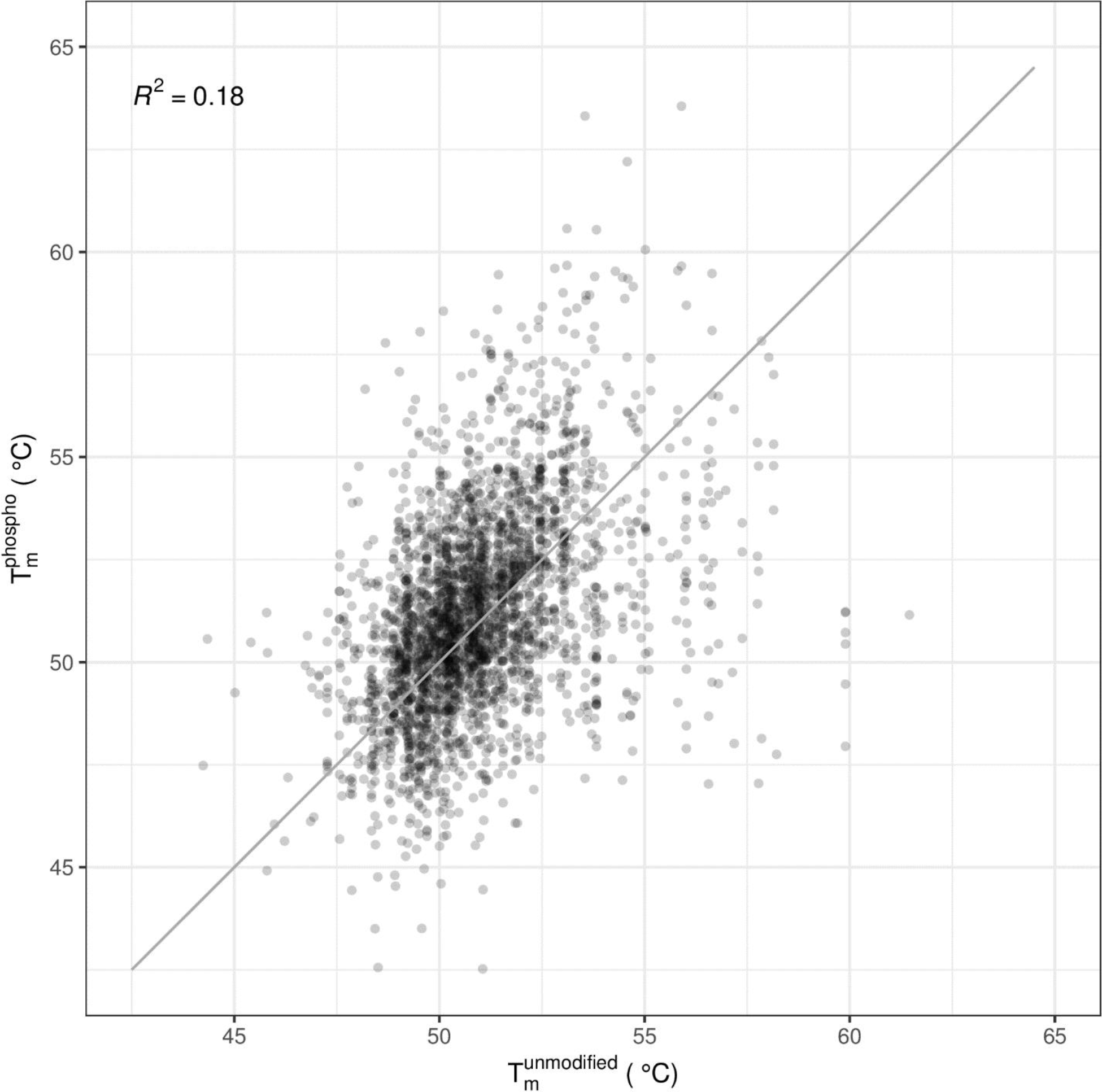
Comparison of non-modified and phosphorylated protein T_m_ estimates from Huang et al. Data collected from supplementary tables 1 and 2 of Huang et al. for high confidence T_m_ estimates and average of all replicates.

**Supplementary figure 2.**
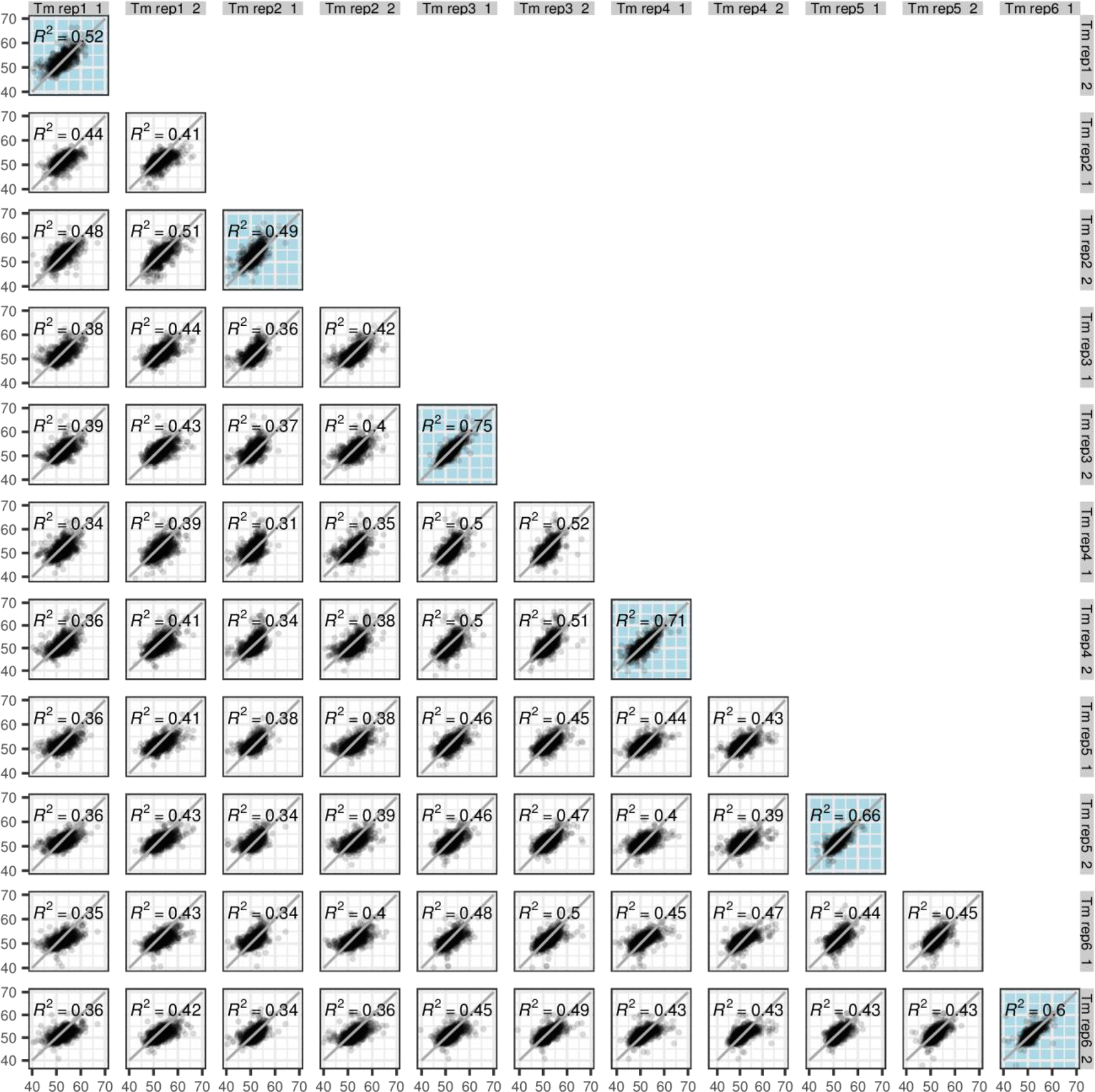
Reproducibility of non-modified sample T_m_ estimates by Huang et al. Displayed T_m_ are represented in degree Celsius, the gray line is the identity line, plots with colored background represent technical replicates according to the labelling by Huang et al. Of note, the data could not be filtered according to the high quality criteria used by Huang et al. because this information was not annotated in the supplementary table supplied.

**Supplementary figure 3.**
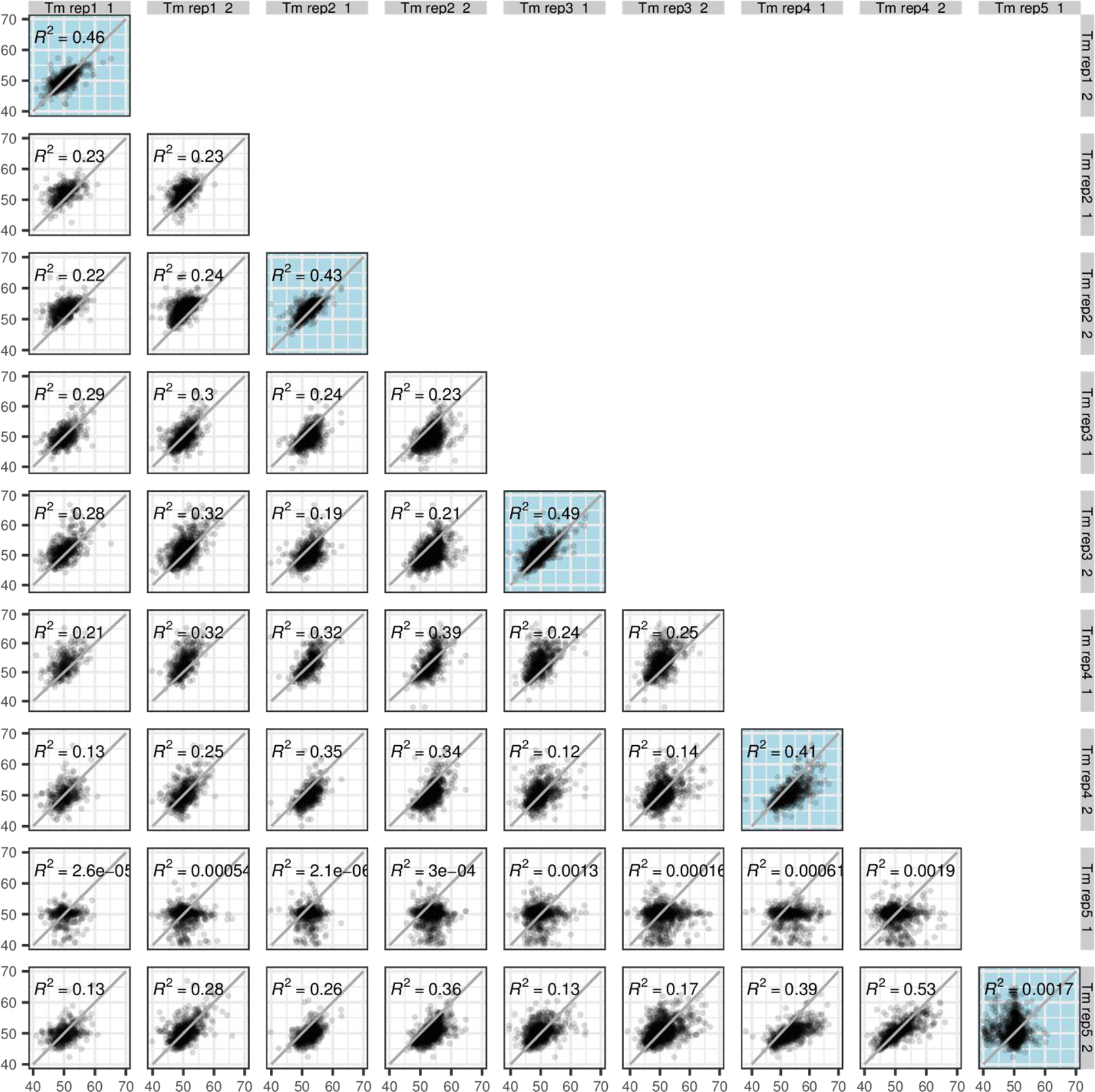
Reproducibility of high confidence phosphorylated protein T_m_ estimates by Huang et al. Displayed Tm are represented in degree Celsius, the gray line is the identity line, plots with colored background represent technical replicates according to the labelling by Huang et al.

**Supplementary figure 4.**
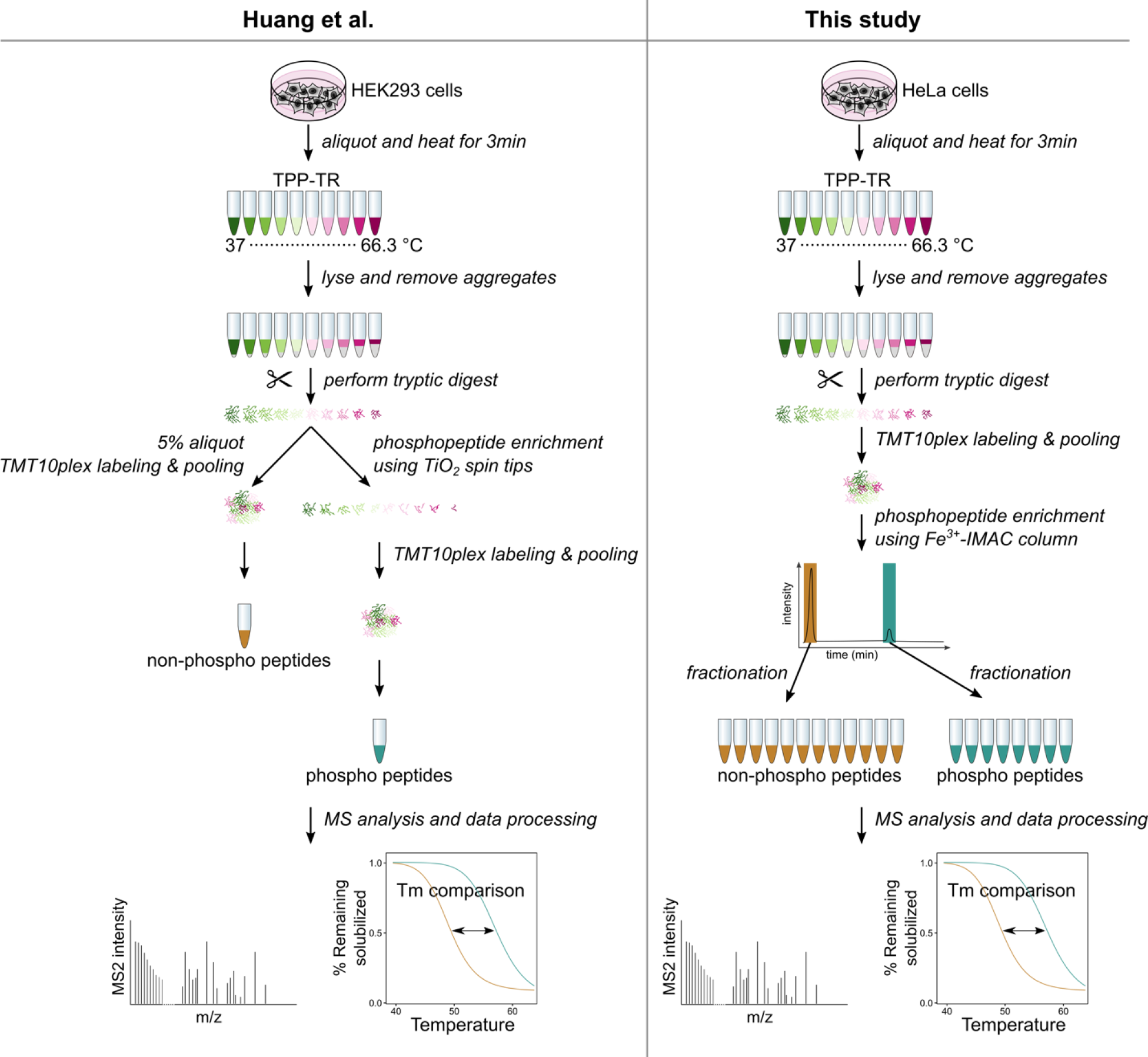
Comparison of workflow from Huang et al. and that of this study. In this study, HeLa cells were heated to ten different temperatures. After lysis, the remaining soluble proteins were collected, digested and TMT labeled. Phosphopeptides were enriched using an iron ion-immobilized metal affinity chromatography (Fe^3+^-IMAC) column and fractionated onto six fractions. The flow-through of the enrichment (non-modified) was fractionated onto twelve fractions. All samples were analyzed on a Fusion Lumos Orbitrap coupled to liquid chromatography.

**Supplementary figure 5.**
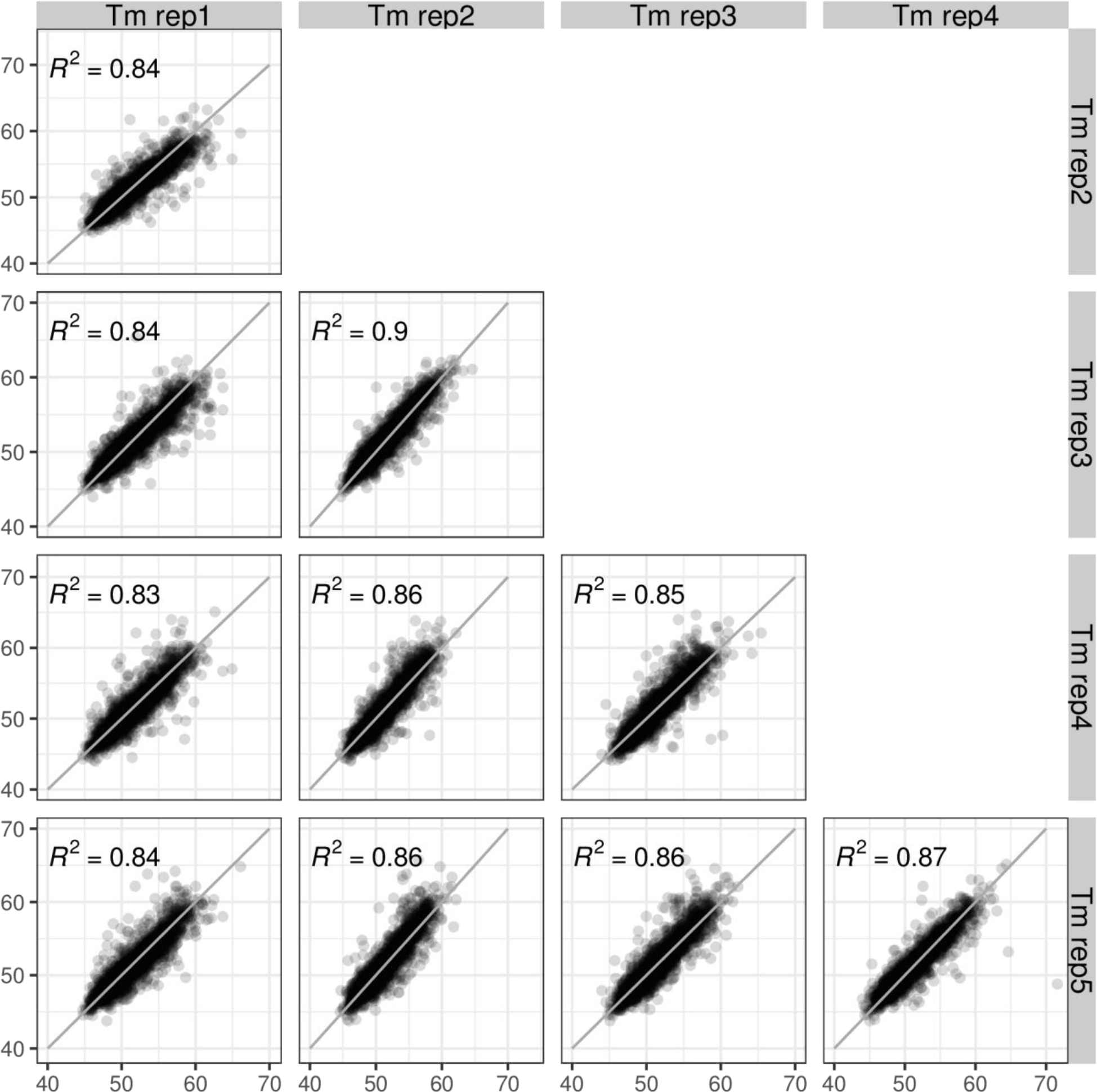
Reproducibility of non-modified sample T_m_ estimates from this study. Displayed T_m_ are represented in degree Celsius, the gray line is the identity line.

**Supplementary figure 6.**
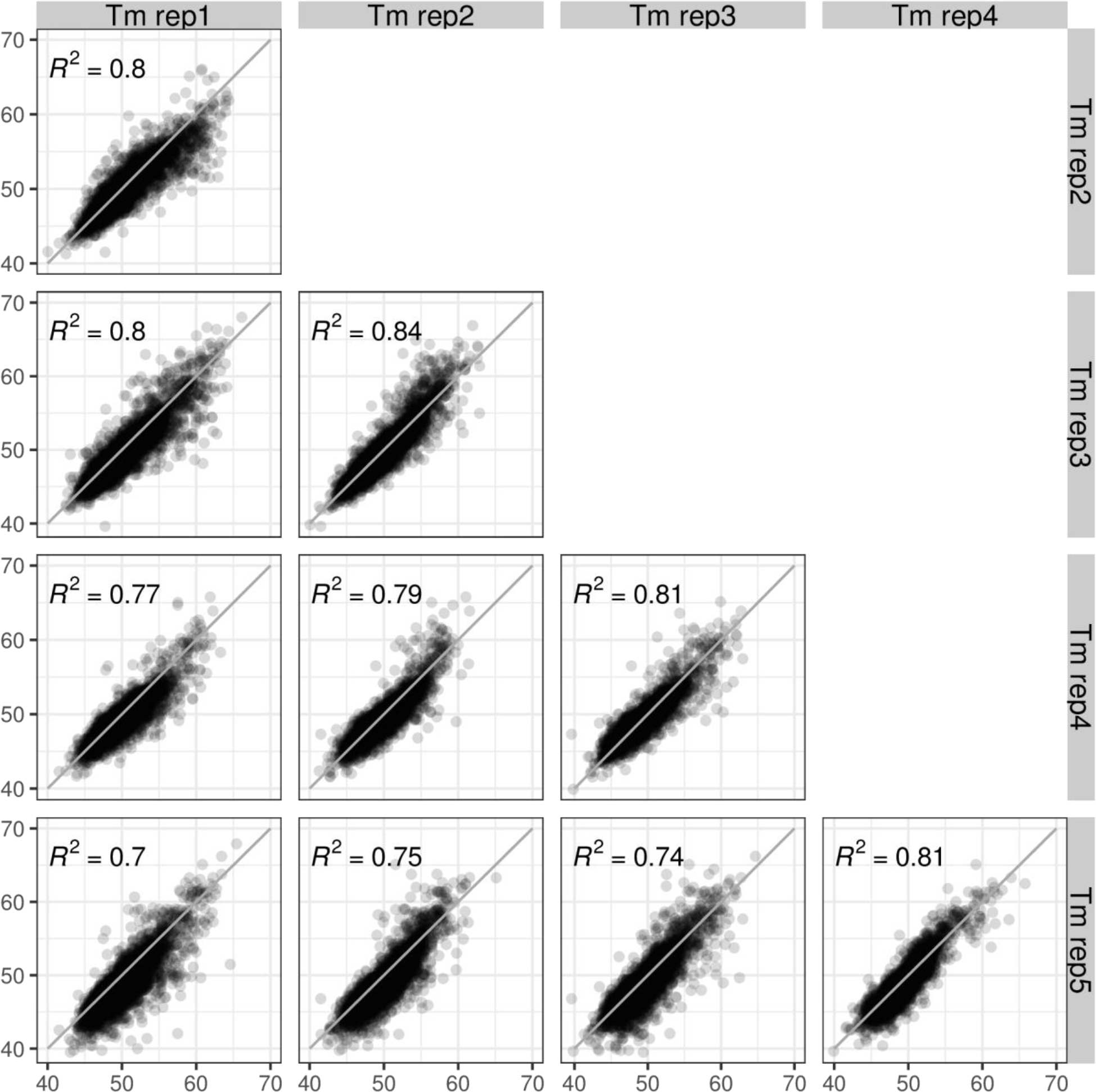
Reproducibility of phosphorylated protein T_m_ estimates from this study. Displayed T_m_ are represented in degree Celsius, the gray line is the identity line.

**Supplementary figure 7.**
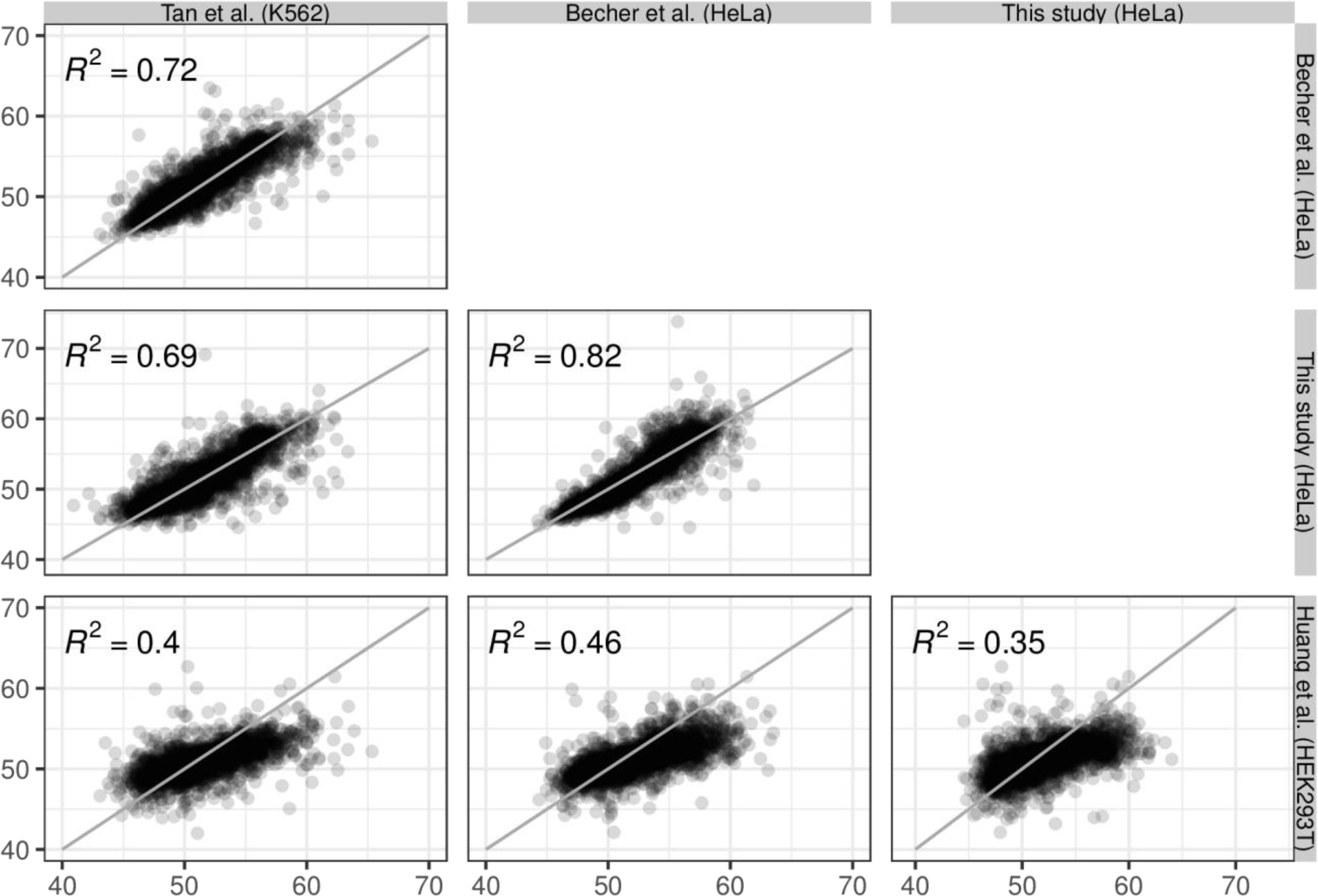
Comparison of T_m_ estimates from Huang et al and this study with previously published datasets. Displayed T_m_ are represented in degree Celsius, the gray line is the identity line. The data from Tan et al.^21^ was retrieved from supplementary table S7 of their publication (K562 cells; using similar acquisition parameters as our work) and fitted using the procedure described in this study. Data from Becher et al.^14^ was retrieved directly from supplementary table S4 (HeLa cells in G1/S phase).

**Supplementary figure 8.**
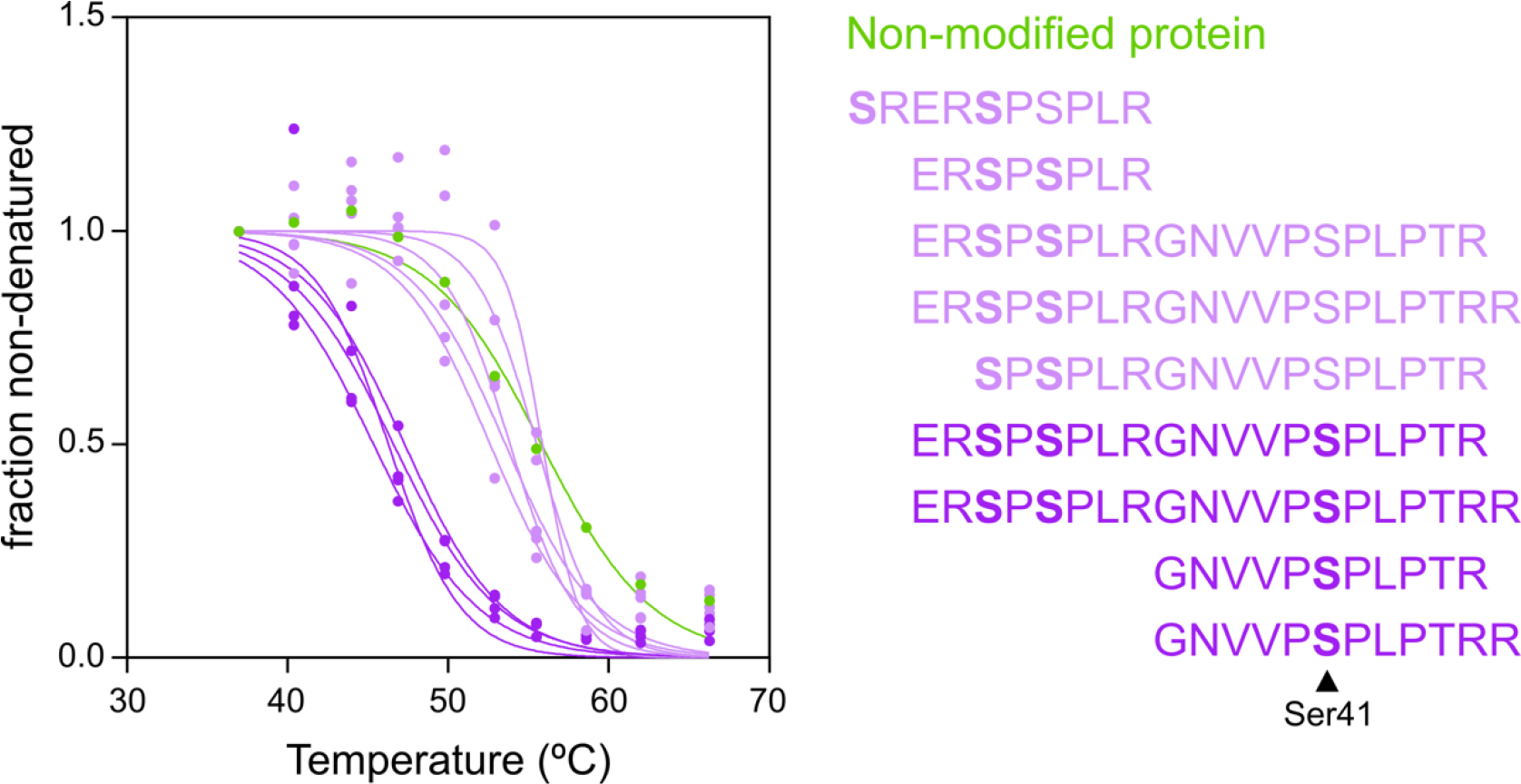
Melting curves of phosphopeptides and non-modified CARHSP1. Bold in the figure legend indicates phosphorylated site.

**Supplementary figure 9.**
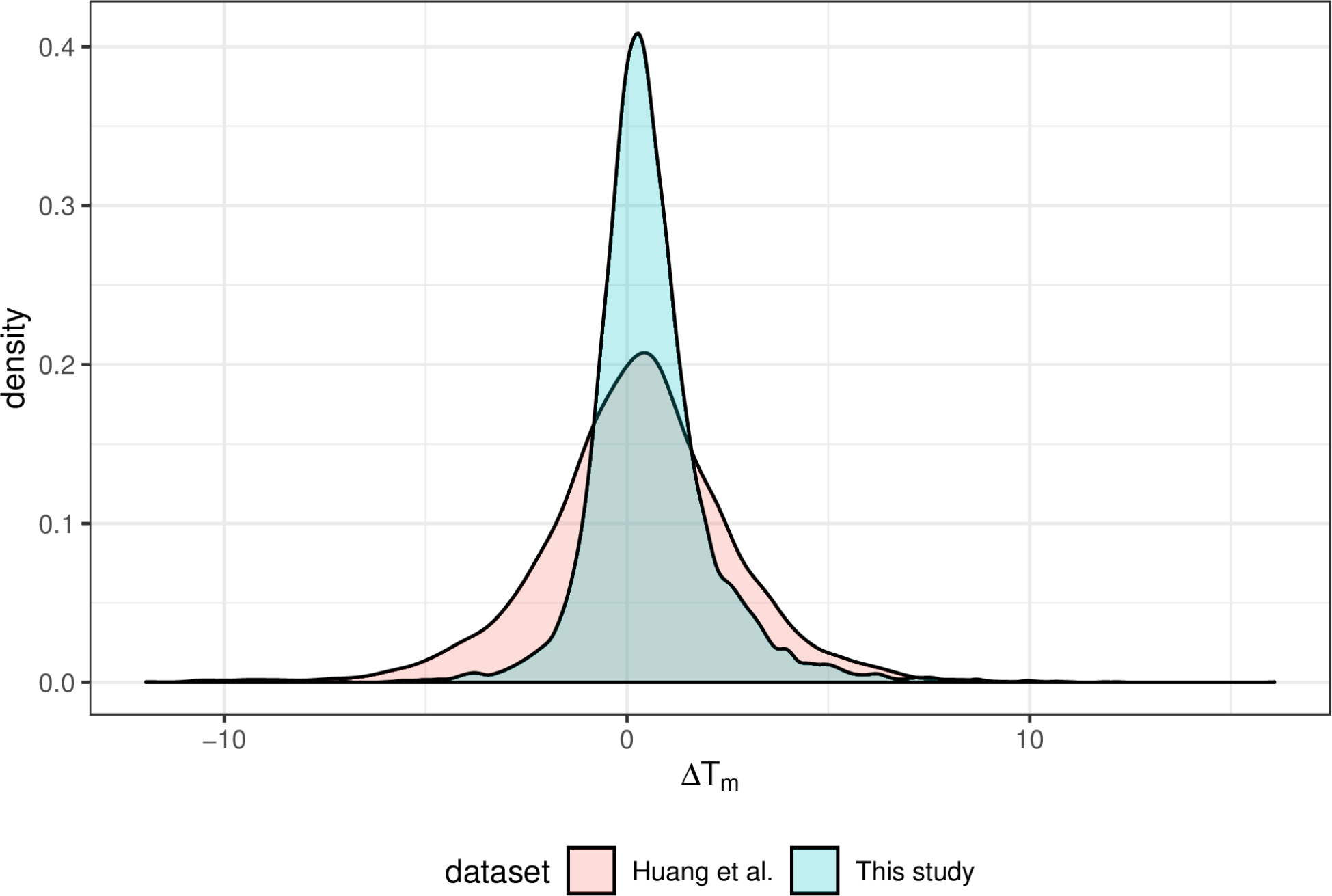
Comparison of ΔTm distribution between the two studies.

**Supplementary figure 10.**
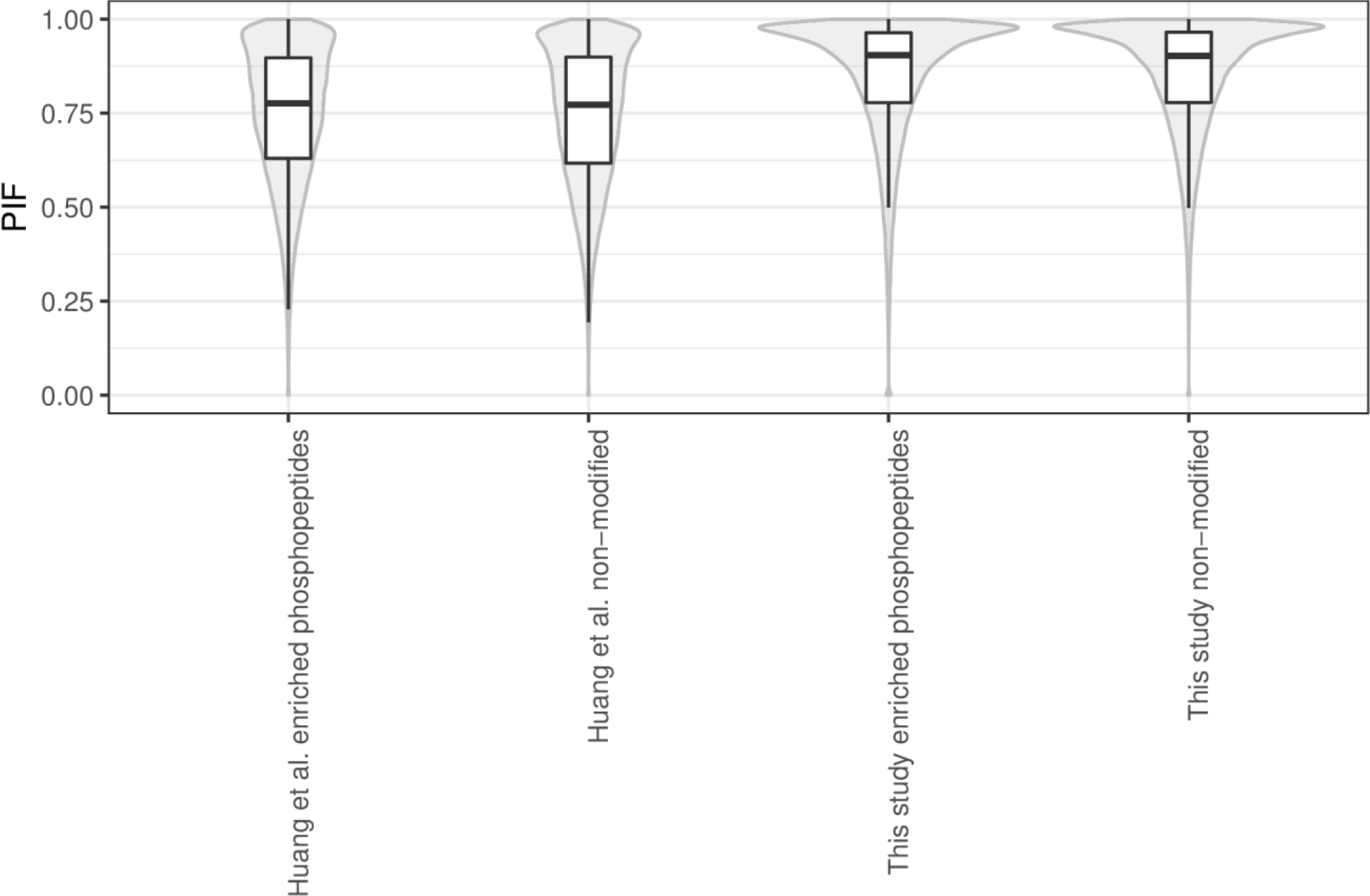
Parent ion fraction (PIF) distribution for the datasets from Huang et al. and this study.

## Supplementary table

**Supplementary table 1.** Protocol differences between the approach used in this study and that in Huang et al.^13^

| **Differences in the protocol presented in this study compared to Huang et al.<sup>13</sup>** | **Reason for change** |
| --- | --- |
| Peptide labeling with tandem mass tags (TMT) prior to phosphopeptide enrichment | To avoid two rounds of TMT labeling and to ensure that phosphopeptide enrichment does not bias the analysis due to different peptide input from samples from different temperatures |
| Peptide pre-fractionation prior to mass spectrometry analysis | To avoid peptide co-isolation during MS2 scans, which leads to ratio compression<sup>1</sup>, which leads to a narrower distribution of $T_m$ |
| Data normalization prior to $T_m$ estimation (between phosphopeptide and non-modified protein samples and between different replicates) | To ensure that different MS experiments are comparable to each other |
| Statistical analysis with multiple testing correction | To reduce the false positive rate |

